# Deciphering the mechanistic basis for the pathological effect of the Gα_o_ E246K mutation in neurodevelopmental disorder

**DOI:** 10.1101/2025.09.03.673757

**Authors:** Isra Sadiya, Irina Nekrasova, Meirav Avital-Shacham, Naomi van Wijk, Keren Zohar, Nir Kalisman, Dina Shneidman-Duhovny, Ehud Banne, Andreea Nissenkorn, Lubov Blumkin, Michal Linial, Mickey Kosloff

## Abstract

Mutations in the GNAO1 gene, which encodes for Gα_o_, a major neuronal G protein, are associated with neurodevelopmental disorders, epilepsy, and movement disorders. We identified and characterized a spontaneous heterozygous GNAO1 E246K mutation in an Israeli female infant with complex developmental delays and substantial motor difficulties. This mutation has been reported in other cases as a prevalent pathogenic mutation in patients with motor dysfunction and a broad range of neurological outcomes. To investigate the molecular and functional consequences of the Gα_o_ E246K mutation, we employed structural modeling and analysis, biochemical assays, mass spectrometry-based proteomics, and cellular functional assays. We show that this mutation does not affect nucleotide binding, nor basal or RGS- accelerated GTP hydrolysis. Despite the E246 position located within a predicted effector binding region, proteomics analysis did not identify any new cellular partners. Instead, we demonstrate that the E246K mutation disrupts the Gα_o_ regulatory GTPase cycle by directly impairing Gβγ dissociation. This impairment overrides the presence of wild-type Gα_o_, explaining the dominant effect of the severe neurogenetic phenotype in the heterozygous background. These findings establish a new molecular mechanism for a GNAO1 mutation with dominant-negative effects on the GTPase regulatory cycle. The insights gained from studying this mechanism of action provide a basis for developing specific and personalized therapeutic strategies based on the outcome of a missense mutation in GNAO1.

## Introduction

Mutations in the GNAO1 gene have been associated with neurological disorders, including epileptic encephalopathy and movement disorders (Nakamura, Kodera et al. 2013, Axeen, Bell et al. 2021, Novelli, Galosi et al. 2023). GNAO1 encodes for Gα_o_, a G protein that constitutes up to 1% of total brain membrane proteins and is crucial for neuronal signaling in the central nervous system (Sternweis and Robishaw 1984, Jiang and Bajpayee 2009). Gα_o_ mediates its function by modulating cAMP levels, ion channels, Rho GTPases, and other pathways that affect membrane trafficking, neurotransmitter release, neuronal polarity, or cytoskeletal remodeling (Feng, Khalil et al. 2018, Schirinzi, Garone et al. 2019, Akamine, Okuzono et al. 2020, Delorme, Giron et al. 2021). Upon activation, Gα_o_ dissociates from Gβγ and either entity can proceed to activate signaling cascades by interacting with downstream “effectors”. G protein effectors are specific to a given G protein and include adenylyl cyclase for Gα_s_, phospholipase Cβ for Gα_q_, or ion channels for Gβγ. Signaling is terminated when Gα hydrolyzes GTP and reassociates with Gβγ, a process accelerated by Regulator of G protein signaling (RGS) proteins. Accordingly, the duration of Gα_o_ activity is determined by RGS proteins that accelerate the intrinsic GTPase reaction that turns off the G protein (Ross and Wilkie 2000). Correlating with the essential role of Gα_o_ in neuronal function and development, mutations in GNAO1 have been linked to epileptic encephalopathy and movement disorders, typically presenting with developmental delay (DD), seizures, and hyperkinetic motor dysfunction (Nakamura, Kodera et al. 2013, Feng, Khalil et al. 2018, Kelly, Park et al. 2019, Schirinzi, Garone et al. 2019). Indeed, knockout studies in mice have demonstrated that Gα_o_ deficiency leads to severe neurological impairments, including tremors, seizures, and early mortality (Jiang and Bajpayee 2009). In humans, over 50 pathogenic GNAO1 variants have been sporadically reported, primarily involving de novo missense mutations, but also a few that disrupt splicing sites. To test the impact of these mutations, 15 mutations were introduced into cells and Gα_o_ expression and its ability to inhibit cAMP via the α2A-adrenergic receptor were monitored (Feng, Khalil et al. 2018). These mutations were classified as loss-of-function (LOF) mutations, mainly associated with epileptic encephalopathy, or gain-of-function (GOF) mutations, which correlate with excessive cAMP inhibition and consequently movement disorders, with or without seizures (Feng, Sjogren et al. 2017). However, recent findings suggest that the impact of these mutations can be more complex and in some cases cannot be strictly categorized as LOF or GOF, as some mutations have been shown to prevent activation by the GPCR, increase nucleotide binding, and decrease hydrolysis rate simultaneously (Muntean, Masuho et al. 2021, Wang, Dao et al. 2022, Knight, Obarow et al. 2023, Lasa-Aranzasti, Larasati et al. 2024, Larasati, Thiel et al. 2025)

Some pathogenic variants have been investigated in previous studies, suggesting mechanistic outcomes that may lead to observed pathologies. Three mutations in the Gα N- and C-termini can reduce interactions with Gβγ, affect nucleotide binding and hydrolysis, or alter association with the membrane and thereby disturb the subcellular localization of Gα_o_ (Solis, Koval et al. 2024). Mutations at seven positions across the Gα_o_ P-loop region, which is adjacent to the nucleotide, can indeed lead to reduced guanine nucleotide affinity, or to impaired Gβγ binding, presumably also by affecting nucleotide interactions or impaired activation by GPCRs (Muntean, Masuho et al. 2021, Dominguez-Carral, Ludlam et al. 2023, Knight, Obarow et al. 2023, Knight, Krumm et al. 2024, Larasati, Thiel et al. 2025). Ten mutations in the switch I and switch II regions, two key Gα regions that change conformation upon G protein activation, can impair nucleotide binding and hydrolysis, reduce RGS or Gβγ binding, or alter GPCR interactions (Muntean, Masuho et al. 2021, Larasati, Savitsky et al. 2022, Dominguez-Carral, Ludlam et al. 2023, Knight, Obarow et al. 2023, Larasati, Solis et al. 2023, Lasa-Aranzasti, Larasati et al. 2024, Ludlam, Soliani et al. 2024, Larasati, Thiel et al. 2025). Four mutations in the Gα switch III region, which also undergoes conformational changes upon G protein activation, have been shown to either reduce GTP hydrolysis rates, disrupt Gβγ release, or significantly increase association with the GPCR (Muntean, Masuho et al. 2021, Dominguez-Carral, Ludlam et al. 2023, Knight, Obarow et al. 2023, Lasa-Aranzasti, Larasati et al. 2024, Larasati, Thiel et al. 2025). Note that the three Gα_o_ switch regions are a central part of the Gα interface with Gβγ and therefore, mutations in these regions are assumed to affect Gβγ release directly through alterations in the protein-protein interface. Mutations further along Gα_o_, in the αG helix region can also impair interactions with the GPCR, reduce Gβγ release, or alter nucleotide binding or hydrolysis (Dominguez-Carral, Ludlam et al. 2023, Knight, Obarow et al. 2023, Larasati, Thiel et al. 2025).

Mutations in an additional Gα_o_ position, E246, can also lead to movement disorders, developmental delays, or hypotonia (Menke, Engelen et al. 2016, Feng, Khalil et al. 2018). This position is located in the Gα α3 helix, which overlaps with the part of a Gα region known to mediate interactions with effectors (Sprang, Chen and Du 2007). Experimental evaluations of the effects of the Gα_o_ E246K mutation have reached disparate conclusions (Muntean, Masuho et al. 2021, Larasati, Savitsky et al. 2022, Knight, Obarow et al. 2023, Solis, Koval et al. 2024, Larasati, Thiel et al. 2025). Measuring cAMP production, Feng *et al*. showed that the Gα_o_ E246K mutant led to constitutive Gα activation and thereby increased inhibition of cAMP production (Feng, Sjogren et al. 2017). Measuring nucleotide binding and hydrolysis across an array of Gα_o_ mutants using fluorescent BODIPY GTPγS and GTP, respectively, showed an increase in binding and a reduction in hydrolysis for the E246K mutant, also suggesting that this mutation leads to constitutive Gα_o_ activation (Solis, Koval et al. 2024). The same study also observed an increase in the E246K mutant binding to the Ric8A protein, which is a known partner for several members of the G_i_ subfamily and is consistent with this particular position being involved in Gα partner interactions. However, other data showed that the Gα_o_ E246K mutant impaired Gβγ release, and therefore reduced G protein activation (Muntean, Masuho et al. 2021, Knight, Obarow et al. 2023), or enhanced Gα_o_ interactions with Gβγ to some degree (Solis, Koval et al. 2024). Additionally, results from immunoprecipitations of GFP-tagged Gα_o_ showed decreased interactions with RGS proteins (Larasati, Savitsky et al. 2022, Solis, Koval et al. 2024, Larasati, Thiel et al. 2025). Taken together, the effects of the E246K mutation and the impact of this mutation on Gα_o_ function remain confusing and unclear.

Here, we applied complementary approaches to investigate the molecular and functional consequences of the Gα_o_ E246K mutation. Our study was motivated by a clinical case of an infant with a spontaneous heterozygous E246K GNAO1 mutation that led to a severe neurogenetic disorder, exhibiting severe communication deficits and developmental delay along with motor dysfunction. Using structural modeling and analysis, biochemical assays, mass spectrometry (MS) proteomics, and an assay designed for quantifying molecular interactions in cells based on bioluminescence resonance energy transfer (BRET), we examined how the E246K mutation can affect Gα_o_ partner interactions, nucleotide binding, GTP hydrolysis, activation and subsequent Gβγ dissociation. Our study contributes to a deeper understanding of G protein-mediated signaling as critical to neurodevelopmental disorders and lays the foundation for developing therapeutic interventions for such disorders (Bennett, Krainer and Cleveland 2019).

## Results

### Clinical outcomes of the Gα_o_ E246K mutation

In recent years, many mutations in the GNAO1 gene have been reported (**Fig. 1**). The encoded Gα_o_ protein has emerged as a critical player in neurodevelopmental disorders, particularly those involving early-onset movement disorders and epilepsy (Feng, Sjogren et al. 2017). Among the reported missense mutations in this gene **(Fig. 1),** a single amino acid change at position 246 (Glu246Lys; E246K) is noteworthy due to its recurrent appearance among pathogenic GNAO1 variants. Interestingly, there are mutations in this position that substitute Glu246 to Lys, Gln, Gly or Val. The particular mutation we focus on here involves a substitution in the GNAO1 gene, where G is replaced by A at position 56,385,308 on chromosome 16 (chr16:56385308 G>A; reference ID rs797044951). This change corresponds to a c.736G>A mutation in the canonical NM_020988.3 transcript, which results in the substitution of the glutamic acid (Glu) with lysine (Lys) at position 246 of the protein (p.Glu246Lys; E246K). Among the 64 pathogenic or likely pathogenic missense mutations in GNAO1 reported in the ClinVar database, four positions (G40, G203, R209, and E246) are designated as hotspots (**Fig. 1)**, with each of these four positions characterized by multiple amino acid substitutions with pathological consequences (Arya, Spaeth et al. 2017). The E246K mutation has been identified in 18 individuals across multiple cases and different populations that include Korea and China. For over half of the reported cases, detailed clinical information is available (Ananth, Robichaux-Viehoever et al. 2016, Saitsu, Fukai et al. 2016, Schorling, Dietel et al. 2017, Kelly, Park et al. 2019, Kim, Shim et al. 2020) (**Table 1**).

**Figure 1:**
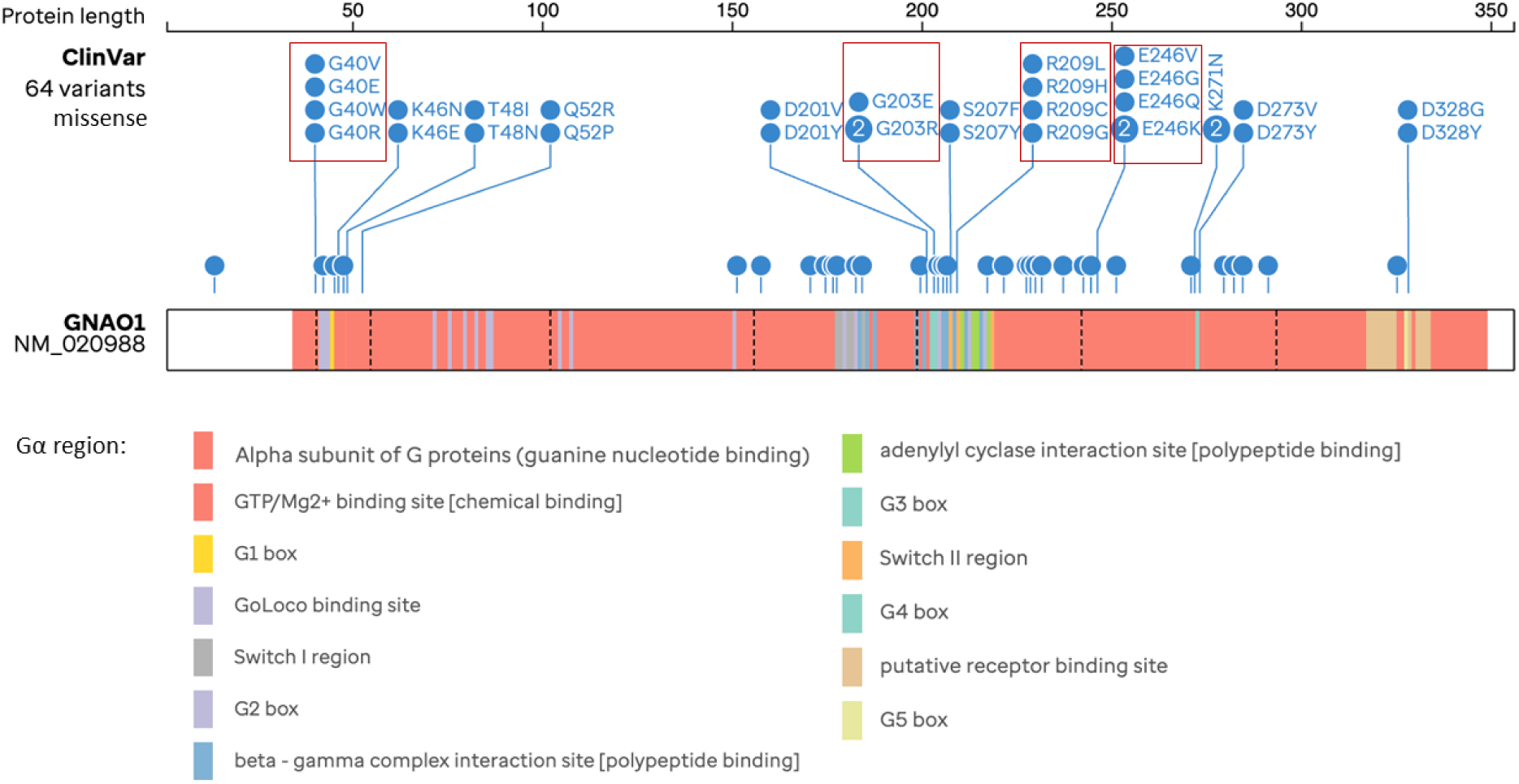
E246 is a hotspot position for mutations in the Gα_o_ protein. Overview of all 64 pathogenic or likely pathogenic missense mutations in GNAO1 (encoding the Gα_o_ protein) reported in ClinVar. Functional domains within the protein of GNAO1 are color coded and labelled. Vertical dashed lines on the protein scheme indicate exon boundaries. Mutations are depicted as blue dots, with their specific positions shown along the protein sequence. Hotspot mutations G40, G203, R209, and E246 are highlighted with red rectangles. These hotspot positions were reported by independent studies, each signified by at least three reports and multiple changes (e.g., G40 changes to V, E, W, R; E246 to K, Q, G, V).

**Table 1.** Clinical outcomes for 13 reported E246K mutation in GNAO1.

| # Cases <sup>a</sup> | Age <sup>b</sup> | Sex <sup>c</sup> | Motor Impairment, DD <sup>d</sup> | Epilepsy | Reference |
| --- | --- | --- | --- | --- | --- |
| 5 | 3m-15m | Mix | Hypotonia; chorea/dystonia by 2 y | 1/5 w focal epilepsy | (Ananth, Robichaux-Viehoever et al. 2016, Saitsu, Fukai et al. 2016) |
| 1 | 7y, 2m | M | Hypotonia, chorea, dystonia | No | (Kim, Shim et al. 2020) |
| 2 <sup>a</sup> | 5y, 6m | M, F | Hypotonia, leads to chorea by ~4 y | No | (Saitsu, Fukai et al. 2016) |
| 2 | 7m; 11m | M, F | DD; variable dystonia | Epilepsy in F only | (Schorling, Dietel et al. 2017) |
| 1 | 3y | F | DD; Hypotonia, variable dystonia | Focal seizures at 7m | (Kelly, Park et al. 2019) |
| 1 | 2y, 3m (d) | M | DD; no prominent movement symptoms | No | (Yang, Niu et al. 2021) |
| 1 | 3y, 9m | F | DD, chorea, dystonia | No | This study |

<sup>a</sup> twin; <sup>b</sup> age by year (y), months (m), death (d); <sup>c</sup> N.D., not determined; <sup>d</sup> DD, developmental delay.

**Table 1** includes cases of E246K mutations along with their reported clinical manifestations (Ananth, Robichaux-Viehoever et al. 2016, Saitsu, Fukai et al. 2016, Kelly, Park et al. 2019). A common phenotype is a developmental delay (DD) and motor impairment (Dzinovic, Skorvanek et al. 2021). In most cases, there is a report on early hypotonia followed by the development of hyperkinetic movement disorders, chorea, dystonia, stereotypies and athetosis, typically between ages 2-4 years. Only 10% of the cases were also reported with seizures and an epilepsy diagnosis (mostly focal-onset). None of the affected individuals had early epileptic encephalopathy. Brain imaging analyses for these patients were inconclusive, as some patients show basal ganglia atrophy or thinning of corpus callosum, while others have normal scans. **Table 1** shows the occurrences of DD and motor impairment and the rare occurrence of epilepsy observed in females. Overall, the data are currently insufficient to generalize whether there is a sex-dependent epileptic trajectory pattern, but we note that most other GNAO1 mutations are associated with severe epileptic encephalopathies.

In this study (**Table 1**) we report on a young Israeli female, diagnosed in 2019 with a neurogenetic disorder at two years of age. Through exome sequencing, confirmed with Sanger sequencing, she was found with a Gα_o_ E246K de-novo mutation. We provide a detailed clinical assessment of physical and developmental changes of this newly reported patient. Note that in many previous reports, clinical descriptions were minimal. Therefore, this expanded assessment should help to link an individual’s genetic background with clinical manifestations. At the age of two months, hypotonia and paucity of spontaneous movements were noted. Axial tone was decreased, while appendicular tone was increased. Feeding difficulties were present, including poor sucking. During the first year of life, hypertonia with scissoring of the lower limbs was observed. After the age of one year, a transition to generalized hypotonia occurred. Motor development was delayed. At six months, the patient was able to attain a high “puppy” position with brief head lifting. She was able to turn her head side-to-side and visually track desired objects. Head control was absent, and right-sided torticollis was noted during the first year (without strabismus). Upper limb use was limited; she used a communication switch, which required significant effort and latency (∼1 minute), often accompanied by involuntary movements during attempted activation. She exhibited generally preserved ocular motor control, with good tracking at home, but intermittent gaze deviation or aversion in unfamiliar environments, particularly when excited. Tongue movements were noted when she was calm and undisturbed (e.g., in the absence of discomfort such as itching or parasitic infection). Language comprehension appeared intact from 6-8 months of age. According to the caregiver, she demonstrated understanding of simple commands and situational cues. For example, she would stick out her tongue when asked, “Where is your tongue?” and touch her nose when asked about it. When upset, she would continue to cry until the caregiver addressed her needs, at which point she would stop and smile or laugh, suggesting preserved intentional communication and emotional responsiveness. Eye contact was appropriate from early infancy. Social smiling was present and consistent. She communicates using crying and laughter. Additional findings included a tendency toward constipation, though with daily bowel movements and no reported urinary difficulties. Appetite was good, but weight gain was suboptimal. She was fed soft and mashed foods without signs of dysphagia or aspiration. Sleep improved markedly with gabapentin, allowing 12 hours of continuous sleep at night. Movement disorder symptoms persisted during daytime. Investigations included normal brain MRI at 9 months, normal BERA at 18 months, and normal EEG at one year. Her last physical examination at the MAGEN center for rare disorders was at 3.9 years of age. The child, seated in a reclined stroller, vocalizes, cries, accepts feeding, and moves the head side-to-side when held; responds to the environment by looking around and watching a video. Physical features include a dolichocephalic head with temporal narrowing, sparse eyebrows, and a tented mouth. Cranial nerve exam is notable for full ocular movements, normal visual tracking, symmetric facial movements, immediate bilateral auditory response, a strong voice, and normal tongue movements. Motor exam shows symmetrical limb range of motion, variable arm tone without spasticity, very low leg tone, normal reflexes, and extensor plantar responses; the child can hold the head vertically and prone for several minutes, attempts reaching for objects, follows video visually, and exhibits intermittent movement abnormalities including possible blepharospasm, tonic eye deviation, choreiform oral movements during feeding, dystonic limb posturing, and dyskinesia/ballismus during excitement. At the age of seven years of age, she suffered from an episode of status dystonicus that required urgent deep-brain stimulation. Since that event, her dystonia has shown relentless progression. She became unable to feed orally, necessitating operative placement of a percutaneous endoscopic gastrostomy tube. Subsequently, she lost all voluntary motor function. Her most recent brain MRI was normal. Currently, at eight years of age, she exhibits profound global neurodevelopmental delay. She is bedridden, cachectic, and entirely dependent for care. Motor development has been completely arrested, with absence of both gross and fine motor abilities, despite only partial symptomatic benefit from treatment.

### Structural modeling and analysis of the Gα_o_ E246K mutation

To understand how the E246K mutation may contribute to neurodevelopmental disorders by potentially disrupting Gα_o_ function or interactions, we analyzed how mutations in this position may alter the structure or interactions of Gα_o_. Since Gα_o_ structures with other partners are limited, we also utilized the Gα_i1_ structure with its high sequence and structure similarity to Gα_o_. We used the X-ray structures of Gα_o_ bound to the GPCR rhodopsin (PDB ID 6FUF), the Gα_i1_βγ heterotrimer (PDB ID 1GP2), and Gα_o_ bound to RGS16 protein (PDB ID 3C7K) (Wall, Coleman et al. 1995, Slep, Kercher et al. 2008, Tsai, Pamula et al. 2018). Visual inspection revealed that E246 is distal to any of these interaction partners **(Fig. 2)**. It is far from the binding site for the GPCR (**Fig. 2A**), and the distance from Gβγ is over 17Å (**Fig. 2B**). E246 is approximately 8Å from RGS16 (**Fig. 2C**), and >6Å from residues in the nucleotide-binding site. Such relatively large distances argue that the position of E246 does not directly influence GTP binding or hydrolysis **(Fig. 2D)**. We further analyzed whether Gα_o_ E246 can affect interactions with protein partners acting in downstream signaling. Since there are no available structural complexes of Gα_o_ with downstream effectors, we extrapolated from the available complexes with downstream effectors of other heterotrimeric Gα subunits (Gα_s_, Gα_q_, Gα_13_, and Gα_i1_) (Tesmer, Kawano et al. 2005, Lutz, Shankaranarayanan et al. 2007, Sammond, Eletr et al. 2007, Chen, Singer et al. 2008, Nishimura, Kitano et al. 2010, Waldo, Ricks et al. 2010, Hajicek, Kukimoto-Niino et al. 2011, Lyon, Begley et al. 2014). We observed that Gα_o_ E246 is located within a predicted effector-binding region that is shared among Gα subunits **(Fig. 2E)**. Refining this interface to include only residues within the previously defined “effector-binding region” (Sprang, Chen and Du 2007), which consists of the α3 helix and its junction with the β5 strand, revealed that Gα_o_ E246 is still within this more restricted predicted Gα_o_ effector-binding region **(Fig. 2F)**. In summary, from a 3D structural perspective, we argue that E246 does not interact directly with GPCRs, Gβγ, RGS proteins, or the nucleotide but is positioned within the predicted effector-binding region of Gα_o_, raising the possibility that the E246K missense mutation contributes to different interactions with potential downstream effectors.

**Figure 2:**
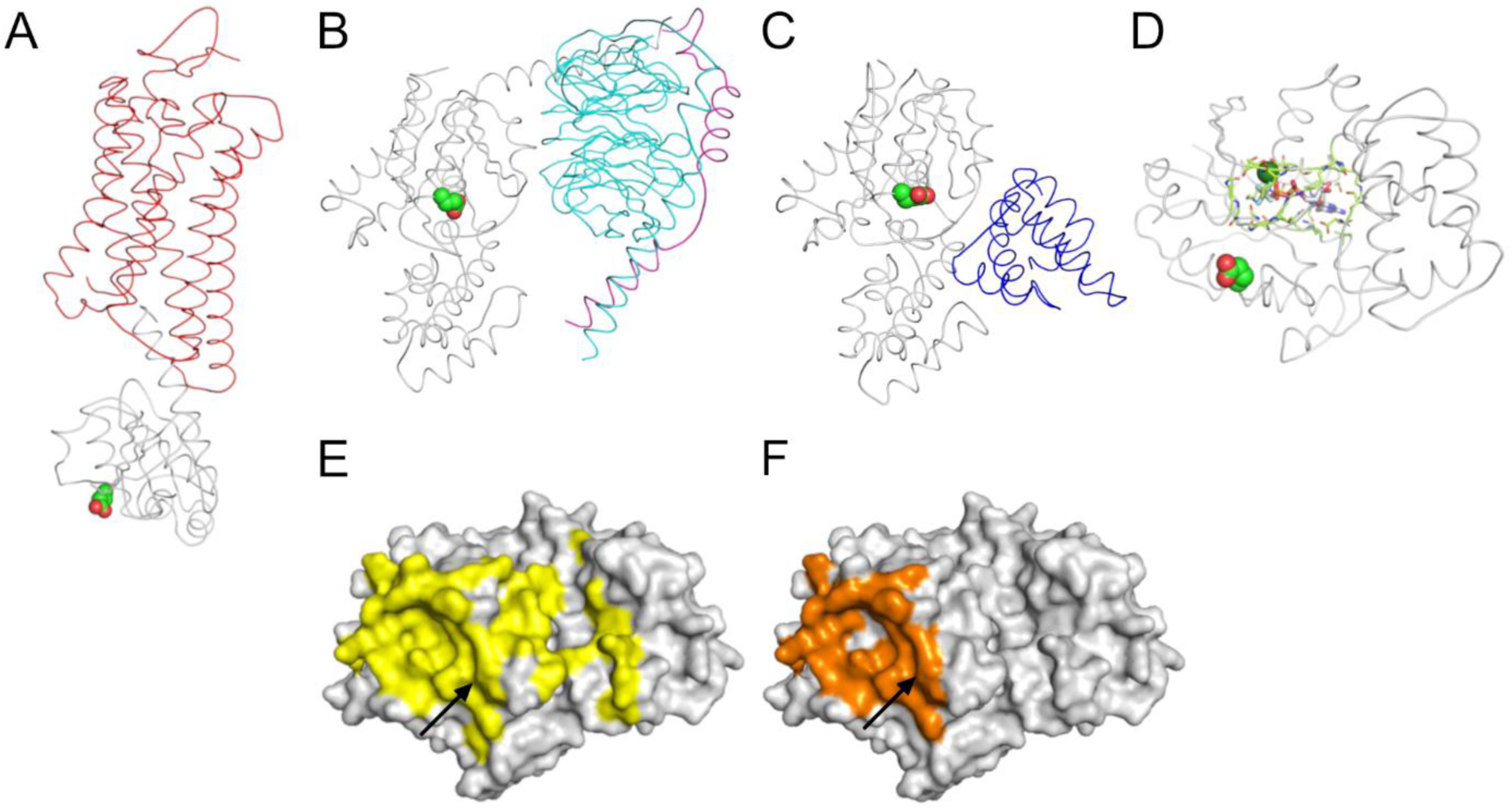
Structural view on Gα_o_ and the E246 position with respect to partner interactions and function. **A**. representative Gα_o_-GPCR (rhodopsin) structure (PDB ID 6FUF) is shown as ribbon diagrams, with Gα_o_ colored gray and the GPCR in red. Gα_o_ E246 is shown as spheres, colored by element. **B**. Gα_i1_ E245 (corresponding to Gα_o_ E246) is > 17Å from Gβγ. The Gαβγ heterotrimer (PDB ID 1GP2) is shown as ribbon diagrams, with Gα_i1_ in gray, Gβ in cyan, and Gγ in magenta**. C**. Gα_o_ E246 is ∼8Å from the RGS protein. The Gα_o_-RGS16 structure (PDB ID 3C7K) is shown as ribbon diagrams, with Gα_o_ in gray and RGS16 in blue. **D**. Gα_o_ E246 is ≥6Å from residues in the nucleotide binding site (defined as all residues ≤5Å from the nucleotide, which are shown as sticks and colored by element). Gα_o_ is shown as a gray ribbon, with the nucleotide analog shown as thick sticks colored by element, and the magnesium ion as a dark green sphere. Gα_o_ is rotated ∼90° about the X axis and ∼135° about the Y axis relative to C. **E**. Gα_o_ E246 is located within the predicted interface of Gα subunits with diverse downstream partners. The Gα_o_ subunit is displayed as in D with a gray molecular surface. The predicted interface is colored yellow and marks positions that participate in Gα interactions with binding partners in at least three of the following structures: Gα_13_–P115RhoGEF (PDB ID 3AB3), Gα_13_–PDZRhoGEF (PDB ID 3CX8), Gα_q_–GRK2 (PDB ID 2BCJ), Gα_q_–p63RhoGEF (PDB ID 2RGN), Gα_q_–PLC-β_3_ (PDB ID 3OHM), Gα_s_–adenylyl cyclase (PDB ID 1AZS), Gα_i1_–βγ (PDB ID 1GP2), and Gα_i1_–RGS14-Goloco (PDB ID 2OM2). The black arrow indicates the position of Gα_o_ E246**. F**. The predicted Gα_o_ “consensus effector-binding region”, colored orange, marking only the subset of positions shown in panel E that are specifically located within the previously-defined “effector-binding region” (Sprang, Chen and Du 2007), extrapolated from the structures of Gα_q_, Gα_13_, Gα_s_, and Gα_i1_ with their respective partners.

### Searching for novel Gα_o_ partners using mass spectrometry

To test whether the Gα_o_ E246K mutant might activate novel effectors or enhance interactions with protein partners (**Fig. 3A**), we performed pull-down experiments from cleared mouse brain homogenates. We added to the homogenates His-tagged wild-type Gα_o_ and Gα_o_ E246K that were recombinantly expressed and purified from *E. coli*, and induced experimental conditions that stabilized active Gα_o_ (see Methods). The interaction of the wild-type and the mutant versions were thus compared in experimental conditions that result in excess activated Gα_o_ and are therefore expected to enhance interactions with any downstream effectors (**Fig. 3A**). We then pulled-down the Gα_o_ proteins by standard His-tag protocol, and analyzed co-eluting proteins by mass spectrometry (**Fig. 3B)**. These experiments showed that the Gα_o_ E246K mutant exhibited slightly reduced interactions with several RGS proteins and increased interaction with Gβγ (Supplementary **Table S1**). Relative to the activated wild-type protein, the activated Gα_o_ E246K mutant showed a slight increase in interactions with the Gβ subunit (Gnb1, **Fig. 3B**), along with decreased interactions with RGS7 and cytoskeletal regulators (e.g., Ywhag and Ywhaz, **Fig 3B**). These altered protein-protein interactions in brain tissues are consistent with disruption in neuronal morphology and synaptic function, concordant with the movement disorder/developmental phenotypes that are seen clinically for E246K carriers – as the Gα_o_ E246K mutant seems to alter neuronal circuits involving motor control. Nevertheless, E246K substitution did not significantly alter effector binding, nor did it enhance binding specificity. Therefore, the MS proteomics results did not show increased interactions with any novel effectors or partners.

**Figure 3:**
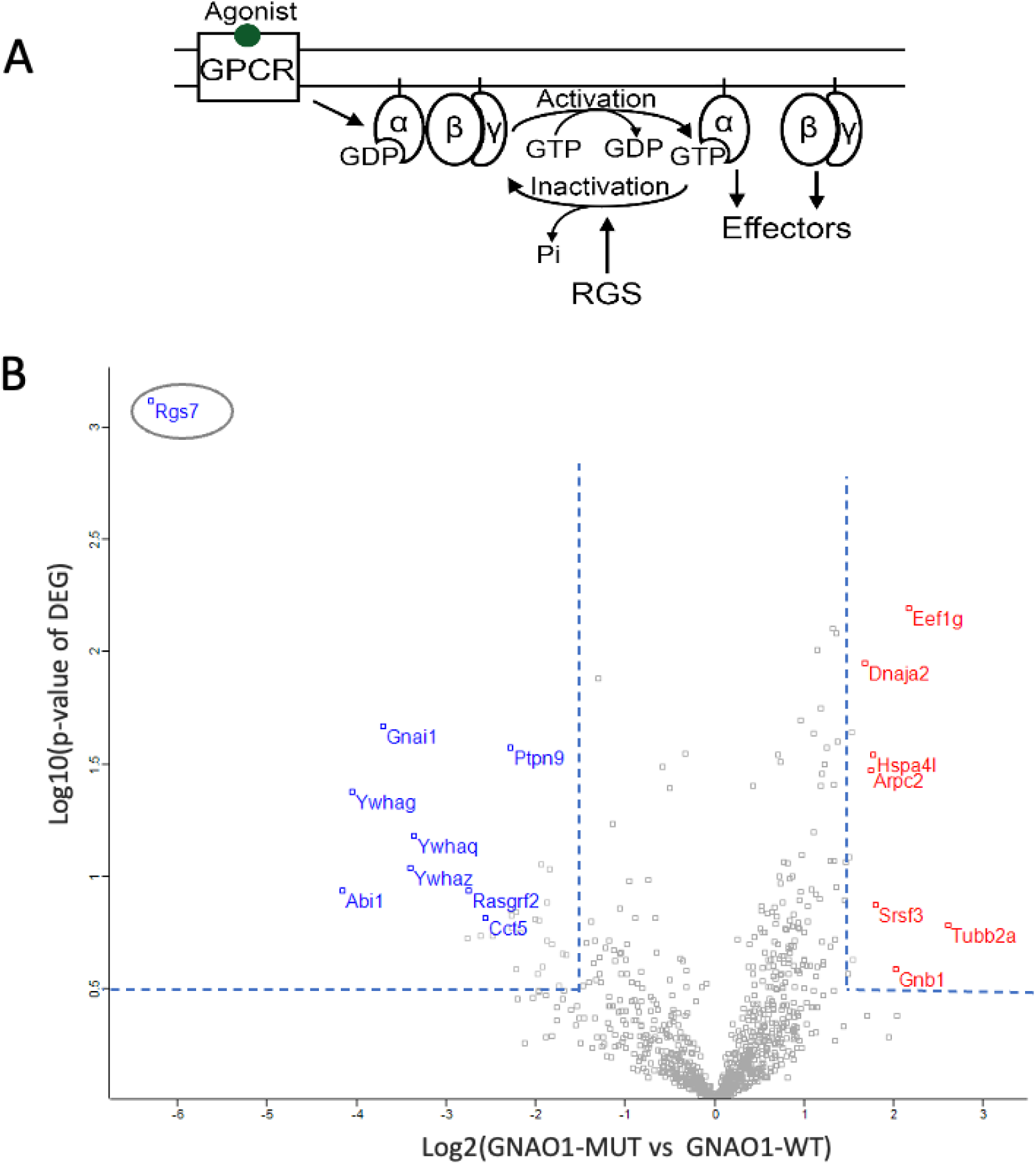
Activated Gα_o_ E246K does not enhance interactions with any novel downstream effectors. **A.** A scheme of the G protein activation cycle. When a ligand activates the GPCR, it induces a conformational change in the receptor that allows the receptor to function as a guanine nucleotide exchange factor (GEF), which exchanges GDP for GTP – thus turning the Gα subunit to an “on” state. This exchange triggers dissociation of Gα-GTP from the Gβγ dimer and from the receptor. Both Gα-GTP and Gβγ can then activate different signaling cascades and effector proteins. RGS proteins act as GTPase Activating Proteins (GAPs), which accelerate the hydrolysis of GTP to GDP by the Gα subunit, thus terminatingthe transduced signal and allowing the Gα subunit to re-associate with Gβγ. **B.** Volcano plot of the results from three biological triplicates following pulldown of the purified Gα proteins. The MS results were collected following an incubation of proteins overexpressed in *E. coli* and purified via nickel-affinity extraction using a His-tag located at the Gα_o_ N-terminus with mouse brain extract. The results compare interactions of the Gα_o_ E246K mutant with wild-type Gα_o_. Dashed lines indicate thresholds for significant fold change in pulldown enrichment (log_2_[DEG] ≥ 1.5, vertical lines) and statistical significance(log_10_[p-value] ≥ 0.5, horizontal lines). Gene symbols mark the most significantly proteins that are underrepresented in the Gα_o_ E246K mutant relative to wild-type, with the top target being RGS7 (circled). Full MS results are provided in Supplementary **Table S1**.

### Biochemical characterization of the Gα_o_ E246K mutant

To determine whether the E246K mutation affects the molecular switch functionalities of Gα_o_, we tested its biochemical properties, including nucleotide binding, basal GTPase activity and its acceleration by RGS proteins. Nucleotide loading was evaluated using an intrinsic tryptophan fluorescence assay with GTPγS. The GTPγS binding rate of Gα_o_ E246K was 0.20 min⁻¹, comparable to that of Gα_o_ wild-type (0.22 min⁻¹). The identical kinetic parameters of GTP binding indicate no effect of the E246K missense variation on nucleotide binding *in vitro* (**Fig. 4A**). Next, we measured GTP hydrolysis rates using a single-turnover GTPase assay. The basal GTPase activity of Gα_o_ E246K was 0.22 ± 0.03 min⁻¹, compared to the rate of wild-type Gα_o_ (0.29 ± 0.01 min⁻¹, **Fig. 4B**). To assess RGS GAP activity, we used RGS16, a representative high-activity RGS domain (Kosloff, Travis et al. 2011). RGS16 exhibited similarly high GAP activity towards both proteins, with a k_GAP_ of 1.6 min⁻¹ for wild-type Gα_o_ and 1.7 min⁻¹ for Gα_o_ E246K (**Fig. 4B**). We also performed dose-response analysis to quantify the GAP potency of RGS16 towards wild-type Gα_o_ and Gα_o_ E246K. Both proteins exhibited similar responses, with comparable EC_50_ values for wild-type Gα_o_ (7 ± 1 nM) and Gα_o_ E246K (10 ± 1 nM); see **Fig. 4C**. Collectively, these results indicate that the E246K mutation does not impair nucleotide binding or GTPase activity or RGS16-mediated GTP hydrolysis.

**Figure 4:**
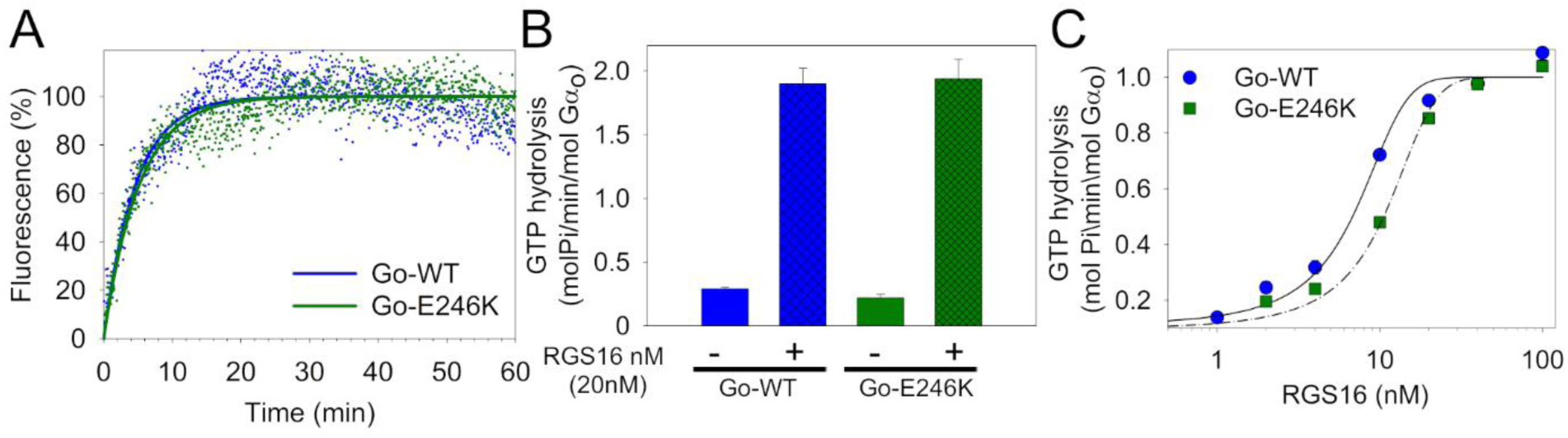
Gα_o_ E246K has no effect on nucleotide binding nor on RGS GAP activity. **A.** GTPγS binding to wild-type Gα_o_ and the Gα_o_ E246K mutant. Nucleotide binding was measured using intrinsic tryptophan fluorescence with excitation at 280 nm and emission measured at 340 nm, initiated by addition of 20 µM GTPγS to 500 nM Gα protein. Binding was measured using relative fluorescence values fitted to a single exponential rise to maximum curve using SigmaPlot 10.0. **B.** GTPase hydrolysis rates with and without RGS16 for Gα_o_ wild-type and the E246K mutant. GTPase rate constants were calculated using SigmaPlot 10.0 from single-exponential fits to the time course of GTP hydrolyzed by the Gα subunits (400 nM), with or without added RGS proteins (20 nM). Values are mean ± s.e.m., n ≥ 3 independent biological replicates. C. Dose responseanalysisof RGS16 activity towards wild-type Gα_o_ and the Gα_o_ E246K mutant. EC_50_ values were calculated using SigmaPlot 10.0 – wild-type Gα_o_ = 7 ± 1 nM, Gα_o_ E246K = 10 ± 1 nM.

We examined the effect of the Gα_o_ E246K mutation on Gα activation and Gβγ dissociation using bioluminescence resonance energy transfer (BRET) in cells (**Fig. 5A**). We used a GPCR kinase-2 (GRK2)-based sensor that measures free Gβγ but does not bind to Gα subunits due to the introduction of the D110A mutation in this protein (Mende, Hundahl et al. 2018). Upon GPCR activation, the GRK2-D110A-GFP sensor was recruited to dissociated Gβγ, causing the GFP conjugated to the sensor to reach the vicinity of the *Renilla luciferase* (Rluc) that is directly conjugated to Gγ. This led to a measurable increase in the BRET signal (**Figs. 5A, 5B**). When we activated the transfected D2R dopamine receptor with 10 μM dopamine, we saw that wild-type Gα_o_ enabled substantial Gβγ release (**Figs. 5B, 5C**).

**Figure 5:**
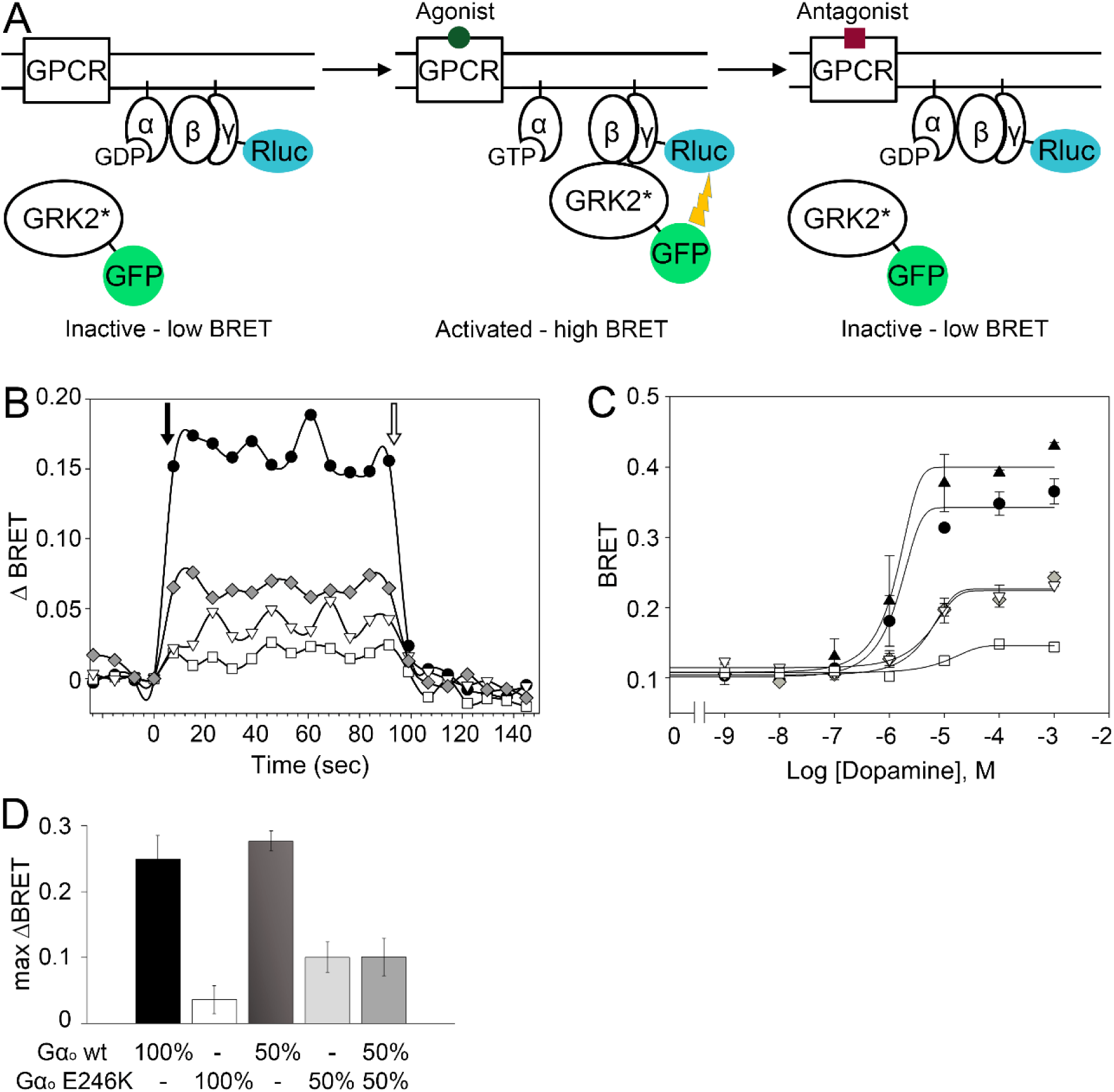
The Gα_o_ E246K mutation decreases Gβγ release upon receptor activation. **A.** Agonist-bound GPCR activates heterotrimeric G proteins, causing the dissociation of the Gα subunit from the Gβγ dimer. The luciferase BRET donor (Rluc, cyan) is fused to the Gβγ dimer, while the fluorescent acceptor (GFP, green) is attached to the GRK protein. Upon activation, free Gβγ-Rluc interacts with GRK-GFP, increasing the BRET signal (ΔBRET). Following antagonist addition, Gα inactivation leads to Gαβγ re-association, decreasing the BRET signal. **B.** Measurements of free Gβγ upon agonist and antagonist additions, with different ratios of wild-type Gα_o_ and the E246K mutant. HEK293 cells were transfected with 20 ng of the D2 dopamine receptor, 10 ng of Gβ_1_, 0.5 ng of Gγ_5_-Rluc, 30 ng of GRK2-D110A-GFP, which binds free Gβγ, and the following ratios of Gα_o_ subunits: 100% wild-type Gα_o_ (30 ng, black circles), 100% Gα_o_ E246K (30 ng, white squares), 50% Gα_o_ E246K (15 ng, white triangles), and 50% wild-type Gα_o_ / 50% Gα_o_ E246K (15 ng each, gray diamonds). Baseline BRET measurements were obtained for 30 seconds before addition of agonist (10 μM dopamine, black arrow). Antagonist (2.5 μM eticlopride) was added following an additional 90 s (white arrow). Each curve is representative of three technical replicates, repeated in three to six biological repetitions. **C.** Dose response analysis of Gβγ dissociation with increasing concentrations of dopamine. Individual experiments were conducted as in B, with data points showing average values ± S.E.M of three technical replicates, repeated in three separate biological repetitions, as follows: 100% wild-type Gα_o_ (30 ng, black circles), 100% Gα_o_ E246K (30 ng, white squares), 50% wild-type Gα_o_ (15 ng, black triangles), 50% Gα_o_ E246K (15 ng, white triangles), and 50% wild-type Gα_o_ / 50% Gα_o_ E246K (15 ng each, gray diamonds). **D.** Maximal ΔBRET values for the experiments shown in C, shown as averages ± S.E.M of three technical repeats, repeated in three separate biological repetitions.

In contrast, the Gα_o_ E246K mutant showed essentially no agonist-dependent increase in the BRET signal, suggesting that this mutation actually prevents Gβγ dissociation. To further test the sensitivity of this function, we repeated the experiment while reducing the amount of transfected Gα_o_ E246K mutant by 50%, which resulted in slightly higher Gβγ release. We conclude that the Gα_o_ E246K mutant inhibits Gβγ dissociation by endogenous Gα subunits. Indeed, co-transfection of the Gα_o_ E246K mutant in equal ratios to wild-type Gα_o_ led to a negligible further increase in Gβγ dissociation at this agonist concentration, which is <30% of the Gβγ released by wild-type Gα_o_ (**Fig. 5B**).

Dose-response analysis with agonist concentrations ranging from 10^-9^ to 10^-3^ M confirmed the inability of the Gα_o_ E246K mutant to mediate Gβγ release upon receptor activation and, importantly, showed the E246K mutation has a dominant suppressive effect over wild-type Gα_o_ (**Figs. 5C, 5D**). Cells overexpressing the Gα_o_ E246K mutant enable release of only ∼15% of the Gβγ that was released by activated wild-type Gα_o_ (**Figs. 5C, 5D**). Transfection of half of the wild-type Gα_o_ led to a slightly higher amount of Gβγ released. Strikingly, a 50%:50% transfection of wild-type Gα_o_ and the Gα_o_ E246K mutant showed ΔBRET responses reaching ∼35% of the response generated by transfecting only the 50% (15 ng) wild-type Gα_o_ – exactly the same extent of the response generated by transfecting only 50% (15 ng) of Gα_o_ E246K. These findings indicate that the Gα_o_ E246K mutation not only significantly impairs Gβγ dissociation, but also exhibits a dominant-negative effect when co-expressed with wild-type Gα_o,_ explaining the clinical phenotype in the de novo heterozygous mutation.

## Discussion

The GNAO1 E246K mutation is recurrently associated with a neurodevelopmental phenotype marked by hypotonia, movement disorders, but mostly without epilepsy. Structurally, E246 resides on the α3 helix of Gα_o_, at a predicted effector-binding interface that is conserved across all 16 Gα subunits. This position is far from the interfaces with the GPCR, Gβγ, and RGS proteins. As the E246 position is also not directly within the nucleotide binding site, our structure analysis suggested the E246K mutation may influence downstream signaling specificity. Despite the structure-based expectation that a mutation in the E246 residue might introduce interactions with novel effectors, we did not identify any such interactions. The only phenotype we observed was decreased interactions with RGS proteins and increased interactions with Gβγ. We also did not observe a significant change in interactions with Ric8A in this assay, as suggested by the results of (Solis, Koval et al. 2024). Furthermore, our findings indicate that the E246K mutation does not impair canonical Gα_o_ functions that include nucleotide binding, GTP hydrolysis or RGS-mediated GAP activity. These biochemical functions of the G protein regulatory cycle showed no significant difference in binding kinetics or EC₅₀ values compared to wild-type Gα_o_. These results challenge earlier hypotheses (Feng, Sjogren et al. 2017, Solis, Koval et al. 2024) that suggested constitutive activation of Gα_o_ by the E246K mutation. Our results do support the suggestions of (Muntean, Masuho et al. 2021, Knight, Obarow et al. 2023), but show the E246K mutation leads to a loss-of-function in regards to Gα_o_-dependent signaling, while some of these previous studies reported this mutant leads to a gain-of-function. We note that our biochemical measurements, unlike previous studies (Larasati, Savitsky et al. 2022, Lasa-Aranzasti, Larasati et al. 2024, Solis, Koval et al. 2024, Larasati, Thiel et al. 2025), did not involve any modifications to the G protein itself. This might explain some apparent discrepancies with previous studies.

Our results present a mechanism of action of the Gα_o_ E246K mutation, that explain how a heterologous mutation leads to the clinical manifestations. We argue that the E246K mutation does not comply with a classic G protein gain-of-function (GOF) mechanism. The functional BRET assays reveal a dramatic reduction in signaling. Upon receptor stimulation, the E246K mutant fails to enable Gβγ release, an essential step in propagating GPCR signaling. This defect appears to be a dominant-negative effect. Co-expression of the E246K variant with wild-type Gα_o_ significantly inhibited Gβγ release, suggesting that the mutant protein blocks the function of wild-type Gα_o_ in heterozygous cells, which resembles the patient’s genetic context. This means that the reduced interaction of the Gα_o_ E246K mutant with RGS proteins observed by us using MS proteomics, and in previous studies (Larasati, Savitsky et al. 2022, Solis, Koval et al. 2024), is not due to reduced affinity, but rather sequestration of Gα_o_ by Gβγ. Our results correspond with the results of Knight *et al*., who used Molecular Dynamics simulations and functional assays to suggest that the corresponding residue in Gα_i1_, E245, is part of a triad with Gα_i1_ residues G203 and R208 that anchors the γ-phosphate of GTP, stabilizing the active state (Knight, Ghosh et al. 2021); disruption of this triad was proposed to affect Gα_i1_ dissociation from Gβγ and thereby impair signaling. Furthermore, a previous structural study of Gα_i1_ proposed that the corresponding residue in Gα_i1_, E245, is allosterically coupled to a conformational rearrangement of the P-loop upon binding to GPCR, destabilizing GDP binding and thereby facilitating GDP release (Zhang, Gui et al. 2021). In a cellular context, Gβγ dimers regulate downstream effectors that include ion channels, kinases, and components of the exocytic machinery. Timely and spatially coordinated Gβγ signaling is presumed to be critical for regulating neurotransmitter release but also for receptor desensitization, and may affect synaptic plasticity. While speculative, these physiological processes may underlie the movement disorders in GNAO1 E246K patients.

Our findings can be transformed to a therapeutic effort that may be designed for restoring G protein subunit cycling while bypassing the defective Gβγ release. Our discovered mechanism for the effects of the GNAO1 E246K mutation may explain why conventional antiepileptic drugs provide little benefit for these primarily movement-based symptoms. The use of tetrabenazine, a dopamine-depleting agent, has been identified as a potential targeted therapy for E246K-related movement disorders (Falsaperla, Sortino et al. 2024). A clue to the function of this agent was based on molecular modeling that supported the idea that tetrabenazine could modulate Gα_o_ activity (Falsaperla, Sortino et al. 2024). Small molecules might therefore be designed to facilitate heterotrimer disassembly or mimic the conformational changes normally triggered by receptor activation (**Fig. 3A**). Due to the dominant effect of E246K in the heterologous setting, it is also attractive to propose the use of oligonucleotides tailored to the specific GNAO1 E246K mutation, as our results show that the mutant suppresses wild-type Gα_o_ activity. It was shown that antisense oligonucleotides (ASOs) that selectively suppress the mutant allele of E246K in patient-derived cells, improved cellular functions such as cAMP signaling (Shomer, Mor et al. 2025). A developed GNAO1 E246K mouse model partially mirrors the human condition and responds to the selected ASO, demonstrating the *in vivo* potential (Shomer, Mor et al. 2025). Thus, precision RNA-based treatments could potentially suppress the mutant allele while preserving normal gene function.

## Materials and methods

### Plasmids and protein constructs

The RGS16 domain was expressed in the pLIC-SGC1 vector as N-terminally His_6_-tagged fusion proteins (Addgene). The N-terminally His_6_-tagged Gα_o_ clone was a gift from Vadim Arshavsky (Duke University). Gα_o_ E246K mutant was generated with the QuikChange site-directed mutagenesis kit (QuickChange Lightning, Agilent Technologies, USA) with primers designed using the Primer Design Program (http://www.genomics.agilent.com). Plasmids used to transfect HEK293T/17 cells were follows: Dopamine D2 receptor (D2R, GenBank: NM_000795), Gα_o_ (GenBank: AH002708) and Gβ1 (GenBank: X04526) in pcDNA3.1(+) were purchased from cDNA Resource Center (Bloomsburg University, https://www.cdna.org). Gα_o_ E246K was created via the site-directed mutagenesis kit as above (QuickChange Lightning, Agilent Technologies, USA). GRK-based BRET biosensor, which monitors the competition between Gα and GPCR kinase-2 (GRK2) for interaction with free Gβγ, was a generous gift from Michel Bouvier (University of Montreal). In this sensor a mutant version of G protein-coupled receptor kinase 2 (GRK2-D110A), which binds exclusively to free Gβγ and lacks interactions with Gα, is fused with GFP which serves as the acceptor (Sterne-Marr, Tesmer et al. 2003, Okashah, Wright et al. 2020), while *Renilla reniformis* luciferase fused to Gγ5 (RlucII-Gγ5) serves as the donor. Upon Gβγ dissociation, GRK2-D110A binds to free Gβ1γ5, leading to an increase in ΔBRET.

### Protein expression and purification

Proteins were expressed in *E. coli* BL21 (DE3) cells and grown in 0.5 or 1 liter of LB broth at 37°C for RGS or Gα proteins, respectively, until an OD_600nm_ ≥1.4 was reached. The temperature was then reduced to 15°C and protein expression was induced by addition of 500 or 100 μM isopropyl-D-thiogalactopyranoside for RGS or Gα proteins, respectively. After 16-18 h cells were harvested by centrifugation at 6,000g for 30 min at 4°C, followed by freezing the pellets at -80°C. Bacterial pellets were suspended in Lysis Buffer (50 mM Tris, pH 8.0, 50 mM NaCl, 5 mM MgCl_2_, 5 mM β-mercaptoethanol, protease inhibitor cocktail (Roche), and 0.5 mM PMSF for G proteins only) and the cells were lysed using a Sonics Vibra-Cell sonicator, followed by centrifugation at 24,000g for 30 min at 4°C. The supernatants were equilibrated to 500 mM NaCl and 20 mM imidazole and loaded onto HisTrapFF 1 ml columns (GE Healthcare Life Sciences). The columns were washed with >20 volumes of Wash Buffer (20 mM Tris, pH 8.0, 500 mM NaCl, 20 mM imidazole) at 4°C and the tagged proteins were eluted with Elution Buffer (20 mM Tris, pH 8.0, 500 mM NaCl, 100 mM imidazole, pH 8.0). The eluate was loaded onto a HiLoad 16/600 Superdex 75 PG gel filtration column (GE Healthcare Life Sciences) at 4°C with ≥1.5 volumes of GF Elution Buffer (50 mM Tris, pH 8.0, 50 mM NaCl, 5 mM β-mercaptoethanol, and 1 mM MgCl_2_ added for Gα subunits only). The eluate was dialyzed against a Dialysis Buffer (GF Elution buffer with 40% glycerol v/v). All purified proteins were estimated to be >95% pure, as assessed by SDS-PAGE electrophoresis and Coomassie staining. Protein concentrations were determined by measuring absorption at A_280nm_, using predicted extinction coefficients (ProtParam, Swiss Institute for Bioinformatics) of the sequence of each expressed protein

### Guanine nucleotide-binding assay

Intrinsic tryptophan fluorescence was used to measure guanine nucleotide binding to heterotrimeric G proteins. All experiments were conducted with a Microplate Reader (SPARK 10M, Tecan) using black-bottomed 96-well plates (NUNC). Time-based assays were conducted with excitation and emission wavelengths set at 280 and 340 nm, respectively. Assays were initiated by addition of 10 µM GTPγS to pre-incubated 500 nM Gα protein in assay buffer containing 50 mM Tris PH 7, 8 mM MgCl_2_, and 1 mM DTT. Fluorescence values were recorded every five seconds, and time-dependent curves were calculated using a fit to an exponential rise to maximum equation in SigmaPlot 10.0 and normalized to the fluorescence values at saturation.

### Single-turnover GTPase assay

We used single-turnover GTPase assays to measure the GTPase activity of Gα proteins with or without various RGS proteins (Ross 2002, Kosloff, Travis et al. 2011). Gα_o_ wild-type, and Gα_o_ E246K mutant were incubated in Reaction Buffer (50 mM HEPES, pH 7.5, 0.05% polyoxyethylene (v/v), 5 mM EDTA, 5 mg/ml BSA, 1 mM dithiothreitol) at 25°C for Gα_o_ and the E246K mutant with 1 μM [γ-^33^P] GTP for 15 min, then cooled on ice for 5 min. GTP hydrolysis was initiated by adding MgCl_2_ to raise the free magnesium concentration to 1 mM, along with 100 μM cold GTP (final concentration), with or without RGS proteins, at 4°C. Aliquots were taken at different time points: for wild-type and mutants without RGS proteins at t = 0.5, 1.5, 3, 6, 12, 17, and 22 min, and with RGS proteins at t = 0.33, 0.66, 1, 1.5, 3, 15, and 20 min. Reactions were quenched with 5% charcoal in 50 mM Na_2_H_2_PO_4_ (pH 3), followed by centrifugation at 12,000g for 5 min at room temperature. The supernatants were transferred to 3 ml of liquid scintillation cocktail and analyzed using a Tri-Carb 2810 TR scintillation counter (Perkin Elmer). GTPase rates were determined from single-exponential fits to the time courses (using SigmaPlot version 10.0). KGAP rate constants were calculated by subtracting the basal GTPase rate (without RGS protein) from the GTPase rate measured in the presence of the RGS protein (Ross 2002, Kosloff, Travis et al. 2011).

### RGS dose-response analysis

Gα_o_ wild-type and the Gα_o_ E246K mutant were loaded with 1 μM [γ-^33^P] GTP for 15 min, at 25°C, then cooled on ice. Each assay was initiated by adding 10 μl of RGS protein in different concentrations in assay buffer (1 mM MgCl_2_ and 100 μM cold GTP), to a tube containing 20 μl of Gα subunit (500 nM) on ice. The reactions were terminated after 45 sec by adding 100 μl 5% perchloric acid and quenched with 700 μl 10% (w/v) charcoal slurry in 50 mM phosphate buffer (pH 7.5), followed by centrifugation at 12,000g for 5 min at room temperature. 200 μl of the supernatants were transferred to 3 ml liquid scintillation and analyzed using a Tri-Carb 2810 TR scintillation counter (Perkin Elmer). EC_50_ values were determined using a three-parameter sigmoidal curve using SigmaPlot 10.0.

### Mass spectrometry Gα_o_ interactions

Mouse brain extract was contributed by the Citri lab (Hebrew University, approved by the research animal ethical committee of the University). Following centrifugation, the supernatant was incubated with purified Gα_o_ wild-type or mutant proteins, corresponding to the rat GNAO1 sequence. Binding was performed in the presence of GDP, NaF, and AlCl₃ to promote complex formation with GDP-AlF_3_ in the G protein active site. After a 2-hour incubation at 4°C, samples were subjected to Ni-affinity pulldown using Ni-NTA resin for the His-tagged Gα_o_ subunit. Bound proteins were washed and eluted with increasing amounts of the imidazole-containing buffer. Eluted samples were acetone-precipitated and prepared for mass spectrometry analysis as previously described (Slavin, Zamel et al. 2021).

### Bioluminescence Resonance Energy Transfer (BRET) assay

HEK293T/17 cells obtained from ATTC (Manassas, VA) were maintained in a 37°C incubator (with 5% CO2) in ATCC Dulbecco’s modified Eagle’s medium (DMEM) with 10% fetal bovine serum antibiotics (100 units/ml penicillin and 100 mg/ml streptomycin). Readings were recorded using a Microplate Reader (SPARK 10M, Tecan) with an acceptor filter of 540 ± 35 nm and a donor filter of 400 ± 40 nm. All measurements were performed at 37°C. The cells (at a cell density 350,000 cells per mL) were transfected with D2 dopamine receptor (D2R, 20 ng), Gβ_1_ (10 ng), Gγ_5_-Rluc (0.5 ng), GRK2*(D110A)-GFP10 (30 ng), and varying ratios of wild-type or E246K mutant Gα_o_ subunits (30 ng total: either 30 ng of wild-type, 30 ng of mutant, or 15 ng of each for a 50:50 mix), with these amounts corresponding to each well of a 96-well plate. The total DNA amount was adjusted with salmon sperm DNA as needed. Transfections were performed using a polyJet transfection reagent (SignaGen), and cells were seeded in poly-D-lysine pretreated white 96-well plates (100 μl per well). Forty-eight hours post-transfection, cells were washed with phosphate-buffered saline and incubated with Tyrode buffer (NaCl 137 mM, KCl 0.9 mM, MgCl₂ 1 mM, NaHCO₃ 11.9 mM, NaH₂PO₄ 3.6 mM, HEPES 25 mM, d -glucose 5.5 mM). After a 30 minutes incubation at 37°C, 10 μL of the luciferase substrate (coelenterazine D, Biotium) was added, followed by an additional 5.5-minute incubation at 37°C. Basal BRET signals were measured for 30 seconds, after which 10 μM dopamine (agonist) was applied. In experiments, also measuring inactivation of Gα, 2.5 μM antagonist (eticlopride) was added 60 after agonist addition. The change in BRET (ΔBRET) was calculated as the difference between the BRET signal after agonist addition and the baseline BRET signal before agonist addition. The BRET signal is calculated as the ratio of light emitted by acceptor (GRK2-D110-GFP10) to that emitted by donor (Gγ5-RlucII), reflecting the amount of free Gβγ.

## Acknowledgements

We are deeply grateful to the family of the affected child for their trust and generosity in allowing the use of her clinical and genetic information to advance research in this field. This work was supported by the International Development Research Centre (IDRC), the Israel Science Foundation (ISF), and the Azrieli Foundation (grant 3512/19), and by a grant from the Council for Higher Education through the Data Science Research Center at the University of Haifa. K.Z. is supported by the Clore Foundation fellowship.

---

**Fig. S1.**
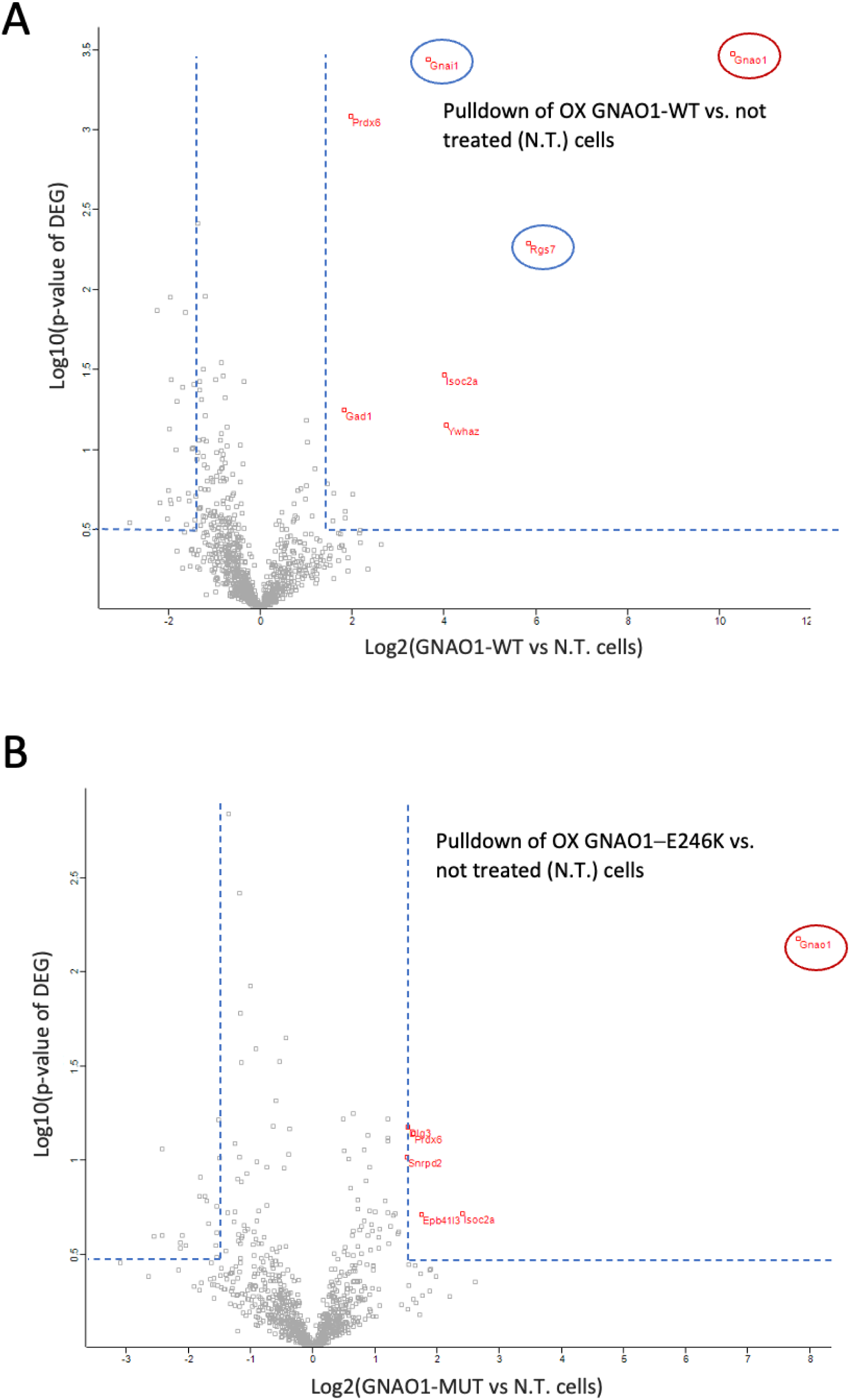
Volcano plot summarizing the results of pulldowns with purified, overexpressed wildtype or the E246K mutant of Gαo incubated with total mouse brain extract. **A.** Comparing treated/untreated wild type Gα_o_ in a pulldown using nickel beads with untreated brain extract (N.T.). As expected, Gα_o_ was the most significantly enriched protein (red circle). Additional enriched proteins included Gα_i1_ and RGS7. RGS7 is a Gα_o_-specific GAP protein that belongs to the RGS family that terminates the G protein signaling cycle. A small number of other enriched proteins, such as Gad1, Prdx6, Iso2a, and a member of the Ywhaz family, co-eluted in the pulldown but are not expected to represent direct interactors of GNAO1. **B**. Comparing treated/untreated Gα_o_ E246K mutant in a pulldown using nickel beads alone with untreated brain extract (N.T.). Each experiment was conducted in triplicate. As expected, Gα_o_, both the wild type and mutant purified overexpressed forms, remained the most enriched protein. Similar associated proteins were identified, though with slightly reduced significance (e.g., Prdx6, Iso2a), along with a few others at borderline significance.

